# EAT-UpTF: Enrichment Analysis Tool for Upstream Transcription Fctors of a gene group

**DOI:** 10.1101/2020.03.22.001537

**Authors:** Sangrea Shim, Pil Joon Seo

## Abstract

EAT-UpTF (<u>E</u>nrichment <u>A</u>nalysis <u>T</u>ool for <u>Up</u>stream <u>T</u>ranscription <u>F</u>actors of a gene group) is an open-source Python script that analyzes the enrichment of upstream transcription factors (TFs) in a group of genes-of-interest (GOIs). EAT-UpTF utilizes genome-wide lists of TF-target genes generated by DNA affinity purification followed by sequencing (DAP-seq) or chromatin immunoprecipitation followed by sequencing (ChIP-seq). Unlike previous methods based on the two-step prediction of *cis*-motifs and DNA-element-binding TFs, our EAT-UpTF analysis enabled a one-step identification of enriched upstream TFs in a set of GOIs using lists of empirically determined TF-target genes.

**Availability:** https://github.com/sangreashim/EAT-UpTF

## INTRODUCTION

The rapid development of high-throughput technologies such as RNA sequencing (RNA-seq), DNA affinity purification followed by sequencing (DAP-seq), and chromatin immunoprecipitation followed by sequencing (ChIP-seq) has led to an explosion in the availability of sequence data. The high-throughput analyses produce lists of genes that are under a particular regulation. When such lists are generated, researchers usually try to understand the biological implications of groups of genes-of-interest (GOIs). To this end, routine follow-up studies typically include gene ontology (GO) enrichment analyses (Maere *et al.*, 2005; Huang *et al.*, 2009) and Kyoto Encyclopedia of Genes and Genomes (KEGG) mapping (Kanehisa and Goto, 2000). In addition, transcription factor (TF) prediction analyses (Kreft *et al.*, 2017; Kulkarni *et al.*, 2018) can be performed to identify consensus upstream regulators of a subset of GOIs, giving a biological insight into the integrated role of the genes under specific conditions. Furthermore, comprehensive identification of TF binding sites and cognate TFs can be used to characterize regulatory networks containing GOIs.

Several bioinformatics tools have been developed to predict consensus DNA sequence motifs in sets of input query genes, especially those from model organisms. For example, *cis*-element sequences that are commonly conserved in the promoter regions of groups of GOIs can be identified using motif enrichment algorithms such as MEME (Bailey *et al.*, 2009). The consensus sequences can then be analyzed using databases of experimentally validated TF binding sites, such as JASPAR (Khan *et al.*, 2018) and TRANSFAC (Matys *et al.*, 2003). However, this two-step approach produces a considerable number of false positives due to over-amplification of error rates caused by double-layered prediction (Kreft *et al.*, 2017). In addition, this method is complicated by the fact that TFs can sometimes bind to gene sequences that differ from their consensus binding sites, and that several TFs undergo protein–protein interactions that enable them to recognize additional DNA sequence motifs. Overall, it is clear that a simplified and realistic prediction of TFs controlling groups of GOIs is necessary to generate a confident conclusion.

O’Malley and colleagues adapted the innovative DAP-seq method and have successfully produced a genome-wide collection of target genes for 349 TFs in *Arabidopsis thaliana* (O’Malley *et al.*, 2016). As a proof-of-concept, we have developed the “<u>E</u>nrichment <u>A</u>nalysis <u>T</u>ool for <u>Up</u>stream <u>T</u>ranscription <u>F</u>actors of a gene group” (EAT-UpTF) tool and combined it with the *Arabidopsis* DAP-seq database to analyze the enrichment of upstream TFs in a group of GOIs. We found that EAT-UpTF was able to robustly evaluate the over-representation of experimentally validated upstream TFs binding to a group of GOIs without the prediction of *cis*-motifs.

## IMPLEMENTATION

High-throughput sequencing analyses typically produce sets of GOIs that require further analyses to evaluate their biological implication and underlying regulatory mechanisms. EAT-UpTF is linked to a DAP-seq database that provides a list of TF-target genes (locus IDs). When a set of GOIs is input in the form of locus IDs, EAT-UpTF identifies the TF targets and compares their relative enrichment in the list of GOIs with that in the total genomic genes. As a result, target genes of certain TFs, which are enriched (over-represented) in the set of GOIs can be identified to predict possible upstream regulators of the gene group. To examine the statistical significance of over-representation, the SciPy module (Oliphant, 2007) is used to perform hypergeometric and binomial tests, which differ in that the binomial test considers replacement whereas the hypergeometric test does not. These two tests are used to compare the occurrence of *x* genes (a subset of TF-target genes) among *n* genes (GOIs) with that of *X* genes (total TF-target genes) among *N* genes (total reference genes). Comparisons with relatively large differences (*x/n – X/N*) can then be considered to identify upstream TFs that may play a role in regulating the set of GOIs.

For the initial validation of EAT-UpTF, we used the DAP-seq *Arabidopsis* database, which lists the target genes of a vast majority of *Arabidopsis* TFs (∼349). Since EAT-UpTF performs enrichment analyses for hundreds of TFs simultaneously, a *post hoc* test should be applied to counteract the type I errors (false positives) originating from multiple testing. A number of *post hoc* analyses can be used to compensate for the increase in the false positive rate caused by multiple tests. The most widely used method is the family-wise error rate (FWER) correction, named after Carlo Emilio Bonferroni. The Bonferroni correction tests individual hypotheses at a significance level of *a*/*m*, where *a* is the desirable alpha level and *m* is the number of tests performed (Bonferroni *et al.*, 1936; Dunn, 1961). This correction method is considered conservative when a large number of tests are conducted, but was likely appropriate in our analysis because the multiple hypothesis tests were limited to several hundred TFs. Another *post hoc* analysis option is the false discovery rate (FDR) correction described by Yoav Benjamini and Yosef Hochberg (Benjamini and Hochberg, 1995). The Benjamini-Hochberg FDR correction tests hypotheses at a significance level of *ka/m*, where *a* is the desirable alpha level, *m* is the number of tests performed, and *k* is the rank of the *p*-value of the hypothesis. These two most popular *post hoc* analyses have been implemented in the current version of EAT-UpTF using the Statsmodels module of Python (Seabold and Perktold, 2010).

To further validate the relevance and accuracy of EAT-UpTF, we input a gene set bound by the LATE ELONGATED HYPOCOTYL (LHY) TF in *Arabidopsis*, which was identified via a ChIP-seq analysis (Adams *et al.*, 2018). EAT-UpTF identified LHY as being an over-represented upstream TF in the test set. Specifically, 71.6% of the input genes were predicted to be bound by LHY (Table 1) and LHY was identified as one of the top five enriched TFs in the test set (Table 1). The mismatch between the EAT-UpTF output and the ChIP-seq data might be related to the fact that DAP-seq is generally more stringent than ChIP-seq. Typically, DAP-seq produces a rigorous gene set and usually identifies a smaller number of TF-target genes than ChIP-seq. Indeed, all of the LHY-target genes identified by DAP-seq were included in the list of LHY-target genes identified by ChIP-seq analysis.

**Table 1.** Summary statistics of the upstream transcription factor (TF) enrichment analysis for the *Arabidopsis* gene set bound by LHY (Adams *et al.*, 2018).

| TF ID (AGI ID) | $x^a$ | $n^b$ | Observed (%) | $X^c$ | $N^d$ | Expected (%) | $p$-value | Corrected $p$-value<sup>e</sup> | Gene symbols | Gene names |
| --- | --- | --- | --- | --- | --- | --- | --- | --- | --- | --- |
| AT5G02840 | 287 | 722 | 39.8 | 4110 | 27206 | 15.1 | $5.84 \times 10^{-60}$ | $2.04 \times 10^{-57}$ | *LCL1* | LHY/CCA1-like 1 |
| AT3G09600 | 426 | 722 | 59.0 | 8276 | 27206 | 30.4 | $2.59 \times 10^{-58}$ | $4.52 \times 10^{-56}$ | *RVE8, LCL5* | LHY-CCA1-LIKE5, REVEILLE 8 |
| AT3G56850 | 275 | 722 | 38.1 | 3936 | 27206 | 14.5 | $6.43 \times 10^{-57}$ | $7.48 \times 10^{-55}$ | *AREB3, DPBF3* | ABA-responsive element binding protein 3 |
| AT2G46270 | 318 | 722 | 44.0 | 5255 | 27206 | 19.3 | $2.09 \times 10^{-53}$ | $1.82 \times 10^{-51}$ | *GBF3* | G-box binding factor 3 |
| AT1G01060 | 517 | 722 | 71.6 | 11896 | 27206 | 43.7 | $3.01 \times 10^{-53}$ | $2.10 \times 10^{-51}$ | *LHY* | LATE ELONGATED HYPOCOTYL |
| AT2G36270 | 274 | 722 | 38.0 | 4188 | 27206 | 15.4 | $8.13 \times 10^{-51}$ | $4.73 \times 10^{-49}$ | *ABI5, AtABI5, GIA1* | GROWTH-INSENSITIVITY TO ABA 1, ABA INSENSITIVE 5 |
| AT3G62420 | 327 | 722 | 45.3 | 5764 | 27206 | 21.2 | $7.54 \times 10^{-49}$ | $3.76 \times 10^{-47}$ | *BZIP53, ATBZIP53* | Basic region/leucine zipper motif 53 |
| AT1G18330 | 619 | 722 | 85.7 | 16878 | 27206 | 62.0 | $3.63 \times 10^{-46}$ | $1.58 \times 10^{-44}$ | *EPR1, RVE7* | EARLY-PHYTOCHROME-RESPONSIVE1, REVEILLE 7 |
| AT5G17300 | 585 | 722 | 81.0 | 15403 | 27206 | 56.6 | $6.78 \times 10^{-45}$ | $2.63 \times 10^{-43}$ | *RVE1* | REVEILLE 1 |
| AT1G32150 | 357 | 722 | 49.4 | 6979 | 27206 | 25.7 | $6.05 \times 10^{-44}$ | $2.11 \times 10^{-42}$ | *bZIP68, AtbZIP68* | Basic region/leucine zipper transcription factor 68 |
| AT4G34590 | 381 | 722 | 52.8 | 7781 | 27206 | 28.6 | $1.94 \times 10^{-43}$ | $6.15 \times 10^{-42}$ | *AtbZIP11, GBF6, BZIP11, ATB2* | Arabidopsis thaliana basic leucine-zipper 11, G-box Binding factor 6 |
| AT5G52660 | 224 | 722 | 31.0 | 3280 | 27206 | 12.1 | $6.20 \times 10^{-43}$ | $1.80 \times 10^{-41}$ | | |
| AT5G15830 | 336 | 722 | 46.5 | 6440 | 27206 | 23.7 | $2.94 \times 10^{-42}$ | $7.88 \times 10^{-41}$ | *bZIP3* | Basic leucine-zipper 3 |
| AT2G18160 | 178 | 722 | 24.7 | 2268 | 27206 | 8.3 | $4.60 \times 10^{-41}$ | $1.15 \times 10^{-39}$ | *GBF5, bZIP2, ATBZIP2, FTM3* | Basic leucine-zipper 2, FLORAL TRANSITION AT THE MERISTEM3, G-BOX BINDING FACTOR 5 |
| AT4G01280 | 339 | 722 | 47.0 | 6654 | 27206 | 24.5 | $1.88 \times 10^{-40}$ | $4.38 \times 10^{-39}$ | | |
| AT3G10113 | 579 | 722 | 80.2 | 15664 | 27206 | 57.6 | $6.91 \times 10^{-39}$ | $1.51 \times 10^{-37}$ | | |
| AT1G45249 | 165 | 722 | 22.9 | 2112 | 27206 | 7.8 | $1.45 \times 10^{-37}$ | $2.98 \times 10^{-36}$ | *ABF2, AREB1, ATAREB1* | ABSCISIC ACID RESPONSIVE ELEMENTS-BINDING PROTEIN 2, abscisic acid responsive elements-binding factor 2 |
| AT3G10800 | 132 | 722 | 18.3 | 1469 | 27206 | 5.4 | $7.86 \times 10^{-36}$ | $1.52 \times 10^{-34}$ | *BZIP28* | NA |
| AT4G36780 | 269 | 722 | 37.3 | 4944 | 27206 | 18.2 | $1.02 \times 10^{-34}$ | $1.88 \times 10^{-33}$ | *BEH2* | BES1/BZR1 homolog 2 |
| AT2G35530 | 137 | 722 | 19.0 | 1630 | 27206 | 6.0 | $3.89 \times 10^{-34}$ | $6.79 \times 10^{-33}$ | *bZIP16* | Basic region/leucine zipper transcription factor 16 |

<sup>a</sup> The number of genes bound by the specific TF in the test set.<sup>b</sup> The number of genes in the test set.<sup>c</sup> The number of genes bound by the specific TF in the reference set.<sup>d</sup> The number of genes in the reference set.<sup>e</sup> The $p$ -value after Bonferroni or Benjamini-Hochberg correction.

The EAT-UpTF analysis requires the input of an experimentally validated genome-wide list of TF-target genes. As mentioned above, we used the DAP-seq *Arabidopsis* database for the initial validation of EAT-UpTF. However, the EAT-UpTF analysis is not limited to the use of DAP-seq data and could also employ ChIP-seq data or any database that provides a list of TF-target genes. In this regard, the EAT-UpTF analysis could be expanded to any species for which DAP-seq, ChIP-seq, or other appropriate sequencing data are available. In the future, a large-scale database integrating DAP-seq and ChIP-seq results would aid the identification of *bona fide* upstream TFs for groups of GOIs. EAT-UpTF is an open platform that can be improved by integrating updated TF databases. To ensure convenience for users, TF regulatory networks of GOIs identified by EAT-UpTF can also be visualized in Cytoscape (Figure 1).

**Figure 1.**
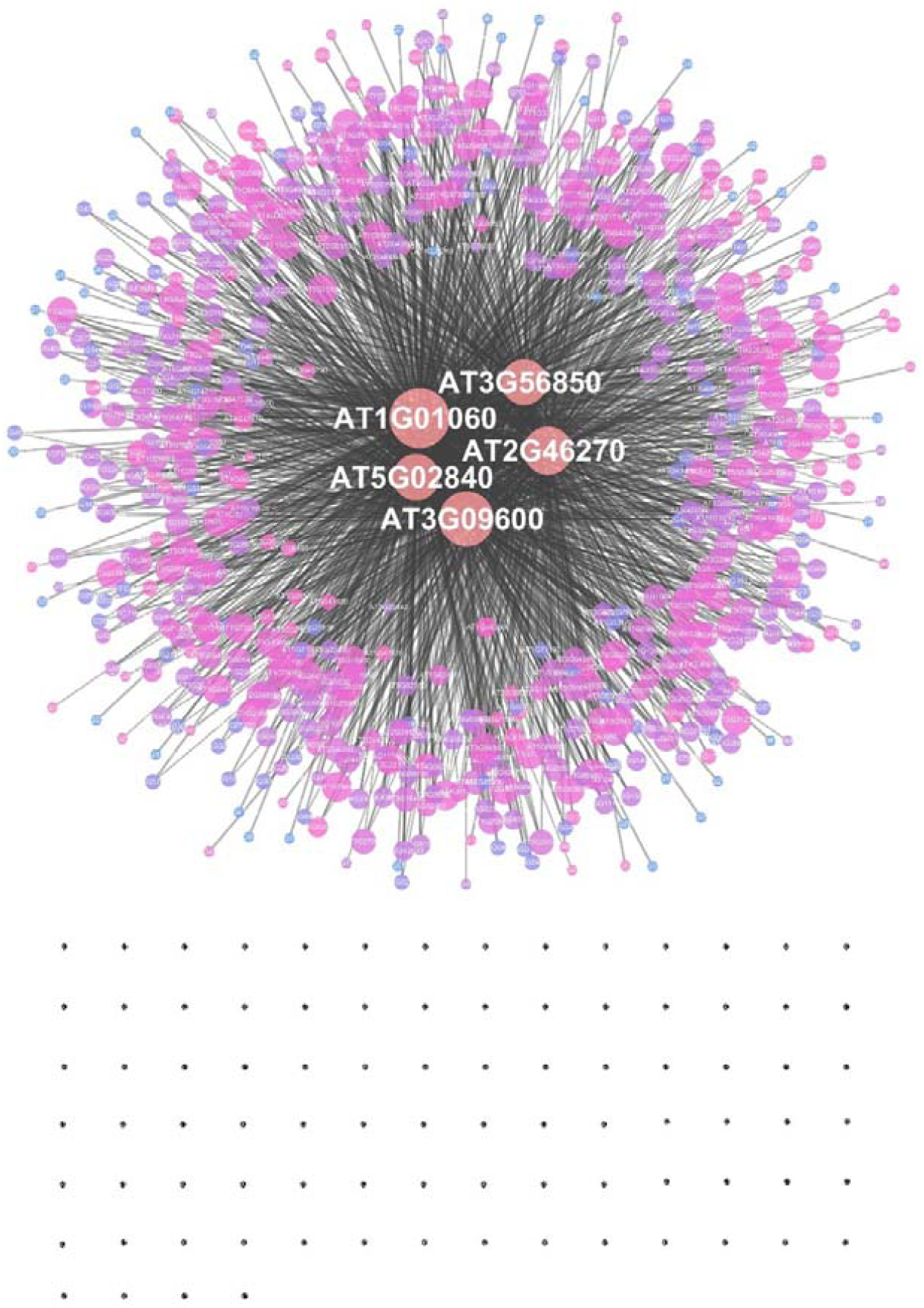
An example of a transcription factor regulatory network constructed by EAT-UpTF. A set of target genes of the LHY transcription factor (Adams *et al.*, 2018) was used as a test input. The area of a node represents the edge count and the color intensity indicates the strength of the neighborhood connectivity. Black dots represent single nodes.

In summary, EAT-UpTF is a tool for analyzing the over-representation of upstream TFs based on the relative enrichment of TF-target genes in a group of GOIs. EAT-UpTF can be used to identify upstream TFs for a group of genes without limitations on species and conservation of *cis*-motifs. With a regular update of databases of TF-target genes, EAT-UpTF will become a powerful tool for TF regulatory network studies.

## Acknowledgments

This work was supported by the National Research Foundation of Korea [NRF-2019R1I1A1A01061376 to S.S, NRF-2019R1A2C2006915 to P.J.S.]; and the Rural Development Administration [PJ01314501 to P.J.S.].

